# Neuronal protein phosphatase 1β regulates glutamate release, cortical myelination, node of Ranvier formation, and action potential propagation in the optic nerve

**DOI:** 10.1101/2024.05.10.593531

**Authors:** Cody McKee, Karl Foley, Katherine M. Andersh, Olivia J. Marola, Brian Wadzinski, Richard T. Libby, Peter Shrager, Houhui Xia

## Abstract

Precise regulation of protein phosphorylation is critical for many cellular processes, and dysfunction in this process has been linked to various neurological disorders and diseases. Protein phosphatase 1 (PP1) is a ubiquitously expressed serine/threonine phosphatase with three major isoforms, (α, β, γ) and hundreds of known substrates. Previously, we reported that PP1α and PP1γ are essential for the known role of PP1 in synaptic physiology and learning/memory, while PP1β displayed a surprising opposing function. *De novo* mutations in PP1β cause neurodevelopmental disorders in humans, but the mechanisms involved are currently unknown. A Cre-Lox system was used to delete PP1β specifically in neurons in order to study its effects on developing mice. These animals fail to survive to 3 postnatal weeks, and exhibit deficits in cortical myelination and glutamate release. There was defective compound action potential (CAP) propagation in the optic nerve of the null mice, which was traced to a deficit in the formation of nodes of Ranvier. Finally, it was found that phosphorylation of the PP1β-specific substrate, myosin light chain 2 (MLC2), is significantly enhanced in PP1β null optic nerves. Several novel important *in vivo* roles of PP1β in neurons were discovered, and these data will aid future investigations in delineating the mechanisms by which *de novo* mutations in PP1β lead to intellectual and developmental delays in patients.

## Introduction

Protein phosphorylation and subsequent dephosphorylation are essential modulatory mechanisms by which distinct cellular processes are regulated. Protein phosphatase 1 (PP1) is a major serine (Ser)/threonine (Thr) phosphatase highly expressed in the brain (Foley et al., 2021). PP1 has been demonstrated to be involved in a myriad of critical cellular processes including transcription, regulation of the cytoskeleton, synaptic transmission, and plasticity (Korrodi-Gregorio et al., 2014). Furthermore, PP1 has been implicated in many pathological processes including Alzheimer’s disease and Noonan syndrome (Braithwaite et al., 2012; Gripp et al., 2016). Additionally, *de novo* mutations in PP1β have been shown to cause intellectual disabilities and developmental delay in human patients (Gripp et al., 2016; Hamdan et al., 2014; Ma et al., 2016). However, given the critical implications of PP1 function, a definitive understanding of the role of PP1 in many of these processes requires further investigation. This is partially due to the existence of three PP1 isoforms, which many studies fail to differentiate among. These three PP1 isoforms are encoded by different genes, with their protein products sharing a similar enzymatic core but different N and C termini, which are important for regulation (Foley et al., 2021). Pharmacological inhibition of PP1, a common and classic way of studying PP1 function *in vivo*, generally does not distinguish among the three isoforms. In order to fully determine the roles of PP1 in various physiological and pathological states it is necessary to understand the relative contributions of the three major PP1 isoforms α, β, and γ in these processes.

By using mice with conditional deletion of the individual PP1 isoforms, we found that PP1γ is the major PP1 isoform in CA1 pyramidal neurons that is critical for synaptic transmission and LTP (Foley et al., 2022; Foley et al., 2023). We previously discovered that PP1β in CA1 pyramidal neurons plays a role opposite to that of PP1γ, in synaptic transmission, LTP induction, and memory formation (Foley et al., 2023). Our work therefore suggests that PP1β plays a previously unknown role in neuronal function. The Cre line used in our earlier study to delete PP1β in a subset of hippocampal neurons begins expressing Cre at ∼P20 (Tsien et al., 1996), and therefore the role of neuronal PP1β function in earlier development was not addressed (Foley et al., 2023).

In the current study, we used a Thy1 promoter to drive Cre expression selectively in neurons. These PP1β cKO mice (Thy1-Cre;*Ppp1cb*^fl/fl^) were born, but exhibited a failure to thrive and typically did not survive past 2-3 postnatal weeks. Using electrophysiology, we determined that PP1β cKO mice have impaired neurotransmitter release at Sch-CA1 synapses. This deficit in neuronal activity coincided with a significant reduction in myelin basic protein (MBP) levels within the cortex. However, we did not observe a change in MBP protein levels in PP1β cKO optic nerves. Interestingly, we detected a robust decrease in the amplitude ratio of the fast:slow components of compound action potentials (CAPs) in PP1β cKO optic nerves. There was no clear difference in the percentage of myelinated axons in these nerves, but we found a significant deficit in the formation of nodes of Ranvier in PP1β cKO optic nerves. Lastly, there was a significant increase in pMLC2 levels in optic nerve axons. Together, these data emphasize the critical role PP1β plays in the development of the CNS including novel roles in influencing *in vivo* neurotransmitter release, cortical myelination, and node of Ranvier formation.

## Methods

### Animals

Thy1-Cre hemizygous mice (JAX #006143) were purchased from the Jackson Laboratory. These mice were then crossed with *Ppp1cb* floxed animals (Liu et al., 2015). Genotypes were validated by PCR using primers for *Ppp1cb* and for Cre-Recombinase. All animals were reared in standard housing conditions and given food/water *ad libitum*. All experiments were conducted using mice at P10-14 (mixed sex).

### Western Blots

Hippocampus and cortex were rapidly dissected and flash frozen on dry ice. Tissues were mechanically homogenized in RIPA (50mM Tris HCl pH 8, 1% NP-40, 0.25% Sodium deoxycholate, 150mM NaCl, 1mM EDTA, 1mM Na_3_VO_4_, 1mM NaF), and protease inhibitor tablets (Thermo scientific A32965) Lysates were then centrifuged at ∼1,000G for 20min at 4°C. Protein concentration was determined using a BCA assay and 20-40μg of protein was loaded per lane. Gels were transferred to PVDF membranes using the Trans-Blot Turbo^®^ Transfer^TM^ System from Biorad. Membranes were washed 3x in TBST prior to blocking in 5% BSA in TBST for 1 hour. Primary antibodies: anti-PP1β (#126 1:1,000 (Strack et al., 1999)), anti-MBP (Biolegend 808401, 1:1,000), Actin (Sigma A2066, 1:4,000), anti-BDNF (Icosagen 327-100, 1:1,000) in 5% BSA TBST were applied to blots. Blots were then gently rocked overnight at 4°C. The following day, primary antibodies were removed and blots were washed 3x with TBST. Azure AC2115/AC2114 HRP conjugated secondary antibodies were then added at 1:10,000 in 5% BSA TBST for 1 hour at room temperature. Blots were quantified using AzureSpot.

### Electrophysiology

#### Paired-Pulse Facilitation (PPF)

300 μm thick acute brain slices were prepared from experimental mice at P12-14 and allowed to recover in artificial cerebrospinal fluid (ASCF) for 1 hour, bubbled with 95% O_2_ / 5% CO_2_ (carbogen). All recordings were conducted under 2-3 mL/min perfusion of ACSF consisting of (in mM) 140 NaCl, 2.5 KCl, 1.3 MgSO_4_, 1.0 NaH_2_PO_4_, 26 NaHCO_3_, 2.5 CaCl_2_, 11.0 D-glucose. Two stimuli were given with varied inter-pulse intervals. The amplitude of the second peak is divided by the amplitude of the first peak to determine the paired-pulse ratio (PPR).

#### Compound action potentials

For complete details see (Shrager & Youngman, 2017). Briefly, mice were euthanized with CO_2_, and optic nerves were dissected and placed in oxygenated artificial cerebral spinal fluid (ACSF) for 1 hour at room temperature. ACSF contained: 125 mM NaCl, 1.25 mM NaH_2_PO_4_, 25 mM D-glucose, 25 mM NaHCO_3_, 2.5 mM CaCl_2_, 1.3 mM MgCl_2_, and 2.5 mM KCl, and was bubbled with 95% O_2_ / 5% CO_2_. Nerves were transferred to a temperature-controlled chamber and perfused with ACSF. Each end of a nerve was drawn into a suction electrode for stimulation (at the retinal end) and recording. Stimuli were applied at 50 µsec duration, and with supramaximal current amplitudes. CAP records were low-pass filtered at 10 kHz and fed into a data processing system for later analysis.

### Electron microscopy

As described in (McKee et al., 2022), optic nerves were immersion fixed in a combination fixative containing 2.0% paraformaldehyde (PFA)/2.5% glutaraldehyde + 0.5% sucrose in 0.1M sodium cacodylate buffer (pH 7.4). After 24 hours of primary fixation the nerves were rinsed in the same buffer and post-fixed for two hours in buffered 1.0% osmium tetroxide/1.5% potassium ferrocyanide, washed in distilled water, dehydrated in a graded series of ethanol to 100% (x3), transitioned into propylene oxide, infiltrated with EPON/Araldite resin overnight, embedded into molds, and polymerized for 48 hours at 60°C. The blocks were sectioned at one micron and stained with Toluidine Blue prior to thin sectioning at 70nm onto slot formvar/carbon coated nickel grids. The grids were stained with aqueous uranyl acetate and lead citrate and examined using a Hitachi 7650 TEM with an attached Gatan 11-megapixel Erlangshen digital camera and Digitalmicrograph software. Images were analyzed with *FIJI,* and *G*-ratios were calculated using an *ImageJ* plug-in (available at http://gratio.efil.de/), from randomly selected axons. Circumferences were manually traced.

### Immunohistochemistry

#### Optic nerve

Optic nerves excised from euthanized mice were placed in 4% PFA for 30min at room temperature. The nerves were then rinsed with PBS before being placed in 15-20% sucrose overnight. The following day nerves were placed in 30% sucrose and stored at 4°C overnight. They were then embedded in Optimal Cutting Temperature (OCT) compound and cryosectioned at 16um. Once dried, sections were placed in PBS for 10min to wash off OCT. Sections stained for nodes were then blocked for 30min in PBTDS (5% donkey serum, 3% BSA, 0.3% Triton X-100 in PBS). Primary antibodies: anti-Pan-NaV (1:1500 see (Rasband et al., 1999)) and anti-Caspr (Abcam Ab34151 1:200) in PBTDS were added to the sections and incubated at 4°C overnight. Primary antibodies were then removed and sections rinsed with PBTDS. Sections were then incubated with Alexa goat anti-mouse 594 and Alexa donkey anti-rabbit 488 (1:400 in PBTDS) for 1 hour at room temperature, protected from light. Sections were rinsed again in PBS, mounted, and allowed to dry overnight prior to confocal imaging. NaV clusters were manually counted by a masked observer using the cell counter tool in *ImageJ*. Average number of images per animal = 14. Sections stained with antibodies against MBP or pMLC2/Pan-NF were placed in Liberate Antibody Binding (L.A.B. Polysciences Inc.), after the initial PBS wash, for 5 min followed by 3x PBS washes. Sections were then permeabilized in 0.5% Triton X-100 in PBS for 30min at 37°C. After 3x PBS washes, sections were blocked in 5% donkey serum in PBS for 1 hour. Sections were then incubated with primary antibodies: anti-MBP (Biolegend 808401, 1:200, anti-pMLC2(S19/T18) (CST 3674, 1:100), anti-Pan-neurofilament (Biolegend 837904, 1:200)) in 5% donkey serum in PBS overnight at 4°C. MBP intensity peaks were measured using the *Plot Profile* function in *ImageJ*. 6 line measurements were made per image with 2-3 images taken per animal. Plot profile data were then analyzed using the *FindPeaks* function in *MATLAB R2021a.* For pMLC2/NF image analysis, pMLC2 and Pan-NF images were automatically thresholded using the Triangle and Default threshold parameters respectively, in *ImageJ*. Measurements of pMLC2 staining area restricted to thresholded area were normalized to NF for each image. Average number of images per animal >16.

#### Retinal flat mounts

Eyes were enucleated and fixed in 4% PFA in 1X PBS at room temperature for 2 hours. For whole mount assessment, retinas were removed from the optic cup and blocked in 10% horse serum and 0.4% TritonX in 1X PBS overnight at 4℃. Retinas were incubated in primary antibody for 72 hours diluted in 10% horse serum, 0.4% TritonX in 1X PBS. Primary antibodies used were rabbit anti-RBPMS (RNA binding protein, mRNA processing factor, GeneTex, 1:500) and mouse anti-TUJ1 (class III beta tubulin, Biolegend, 1:1000). Retinas were then rinsed with 1X PBS prior to secondary antibody incubation overnight at 4℃. Secondary antibodies used were dilated in 1X PBS (Alexa fluor conjugated, Invitrogen). Retinas were washed and mounted ganglion cell layer up in Fluorogel in TRIS buffer (Electron Microscopy Sciences) and allowed to dry at room temperature overnight before storage at 4℃. RBPMS+ TUJ1+ RGCs were quantified following a collection of eight 40x fields per retina. Images were equally spaced 220um from the peripheral edge of the retina with 2 images per quadrant. Image quantifications were manually performed using the cell-counter tool in *ImageJ* by a masked observer across all groups.

### Retinal sections

As mentioned above, eyes were enucleated and fixed in 4% paraformaldehyde in 1X PBS at room temperature for 2 hours. The corneas and lens were removed from each eye and optic cups processed in a sucrose gradient, from 10 to 30 percent in 1X PBS over 5 days at 4℃. Optic cups were then embedded in tissue freezing medium (TFM, VWR) and frozen at -80℃ until sectioning. Optic cups were then cryosectioned at 14um and stored at -20℃ prior to staining. Sections were returned to room temperature on the day of staining and washed with 1X PBS followed by subsequent washes in 0.1% TritonX in 1X PBS. Following initial washes, sections were blocked in 10% horse serum in 0.1%TritonX in 1X PBS for 2 hours at room temperature. Primary antibody diluted in 1xPBS and 0.1%TritonX was placed on the sections overnight at 4℃. Primary antibodies included goat anti-SOX2 (Santa Cruz, 1:250) or goat anti-choline acetyltransferase (ChAT, Millipore, 1:250), rabbit anti-calretinin (Millipore, 1:250), and guinea pig anti-RBPMS (Phosphosolutions, 1:250). Slides were returned to room temperature and rinsed thoroughly in 1X PBS. Secondary antibodies were diluted in 1X PBS and placed on sections for 1 hour incubation. Sections were washed in 1X PBS and placed in DAPI neuronal counter stain solution for 7 minutes. Following a final wash in 1X PBS, slides were coverslipped with Fluorogel in TRIS buffer (EMS) and stored at 4℃ until time of imaging. Sections were imaged at 20x magnification with 3 adjacent sections imaged per animal. As SOX2 and ChAT antibodies were hosted in the same animal (goat), adjacent slides were chosen to ensure at least 3 sections per animal containing each stain were captured. Image quantifications for calretinin+ cells, ChAt+ cells, RBPMS+ RGCs, and SOX2+ cells were manually performed using the cell-counter tool in *ImageJ* by a masked observer across all groups.

### Statistical analyses

Statistical analyses were performed using *GraphPad Prism9.3* software with alpha set to 0.05. Normality was determined using the Shapiro-Wilk test and parametric data were analyzed using two-tailed unpaired t-tests or ANOVA. Non-parametric data were analyzed using a Mann-Whitney U test or Kolmogorov-Smirnov test for frequency distributions.

## Results

To assess how neuronal PP1β influences CNS function we conditionally deleted PP1β by crossing a neuron specific Thy1-Cre mouse line with *Ppp1cb* floxed mice. PP1β cKO pups are viable and were born at frequencies predicted by Mendelian inheritance. These cKO mice appeared to develop normally from birth until ∼P5 when their weight gain and mobility progression began to lag behind that of their littermate controls (Cre negative *Ppp1cb* floxed animals). Western blotting of hippocampus and cortex confirmed a significant depletion of PP1β protein compared to littermate controls (Fig. 1). Work by (Lontay et al., 2012) demonstrated that PP1β is localized to both the pre and postsynaptic terminals. We have shown that deleting PP1β in CA1 pyramidal neurons alters synaptic transmission and LTP induction (Foley et al., 2023). However, in our previous study, PP1β was deleted only in postsynaptic neurons, and therefore the role of PP1β in presynaptic glutamate release in the hippocampus could not be examined. In this study we measured paired-pulse facilitation in Schaffer-Collateral synapses within the hippocampus to assess presynaptic glutamate release probability. PP1β cKO mice exhibited an increase in the paired-pulse ratio (PPR), suggesting a decrease in neurotransmitter release relative to that of littermate controls (Fig. 2).

**Figure 1.**
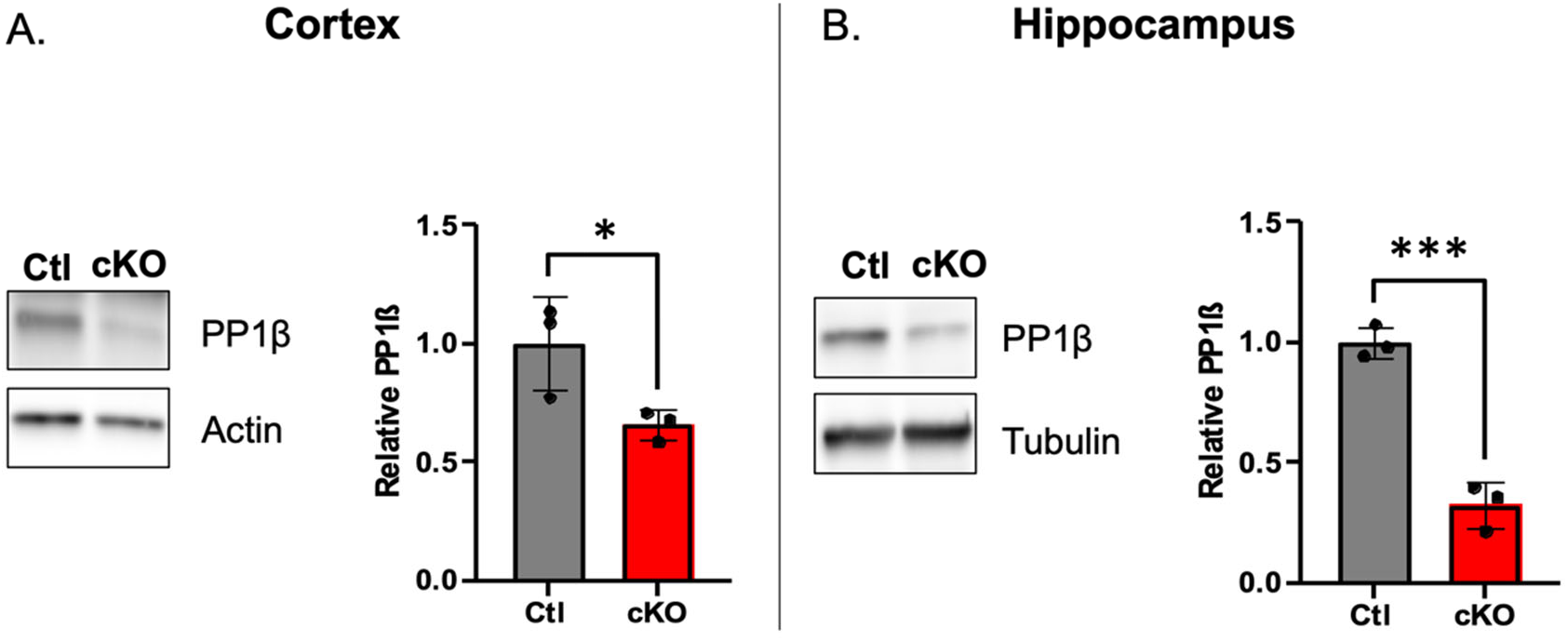
Successful knockdown of PP1β in cortex and hippocampus. **A-B**. Western blot of cortical (**A**) and hippocampal (**B**) tissue from Ctl and PP1β cKO mice probed for PP1β and actin/tubulin (left). Quantification of PP1β normalized to loading control (two-tailed unpaired t-test \**p* < 0.05, \*\*\**p* < 0.001, n = 3 animals, P12; Error bars denote mean +/-SEM) (right).

**Figure 2.**
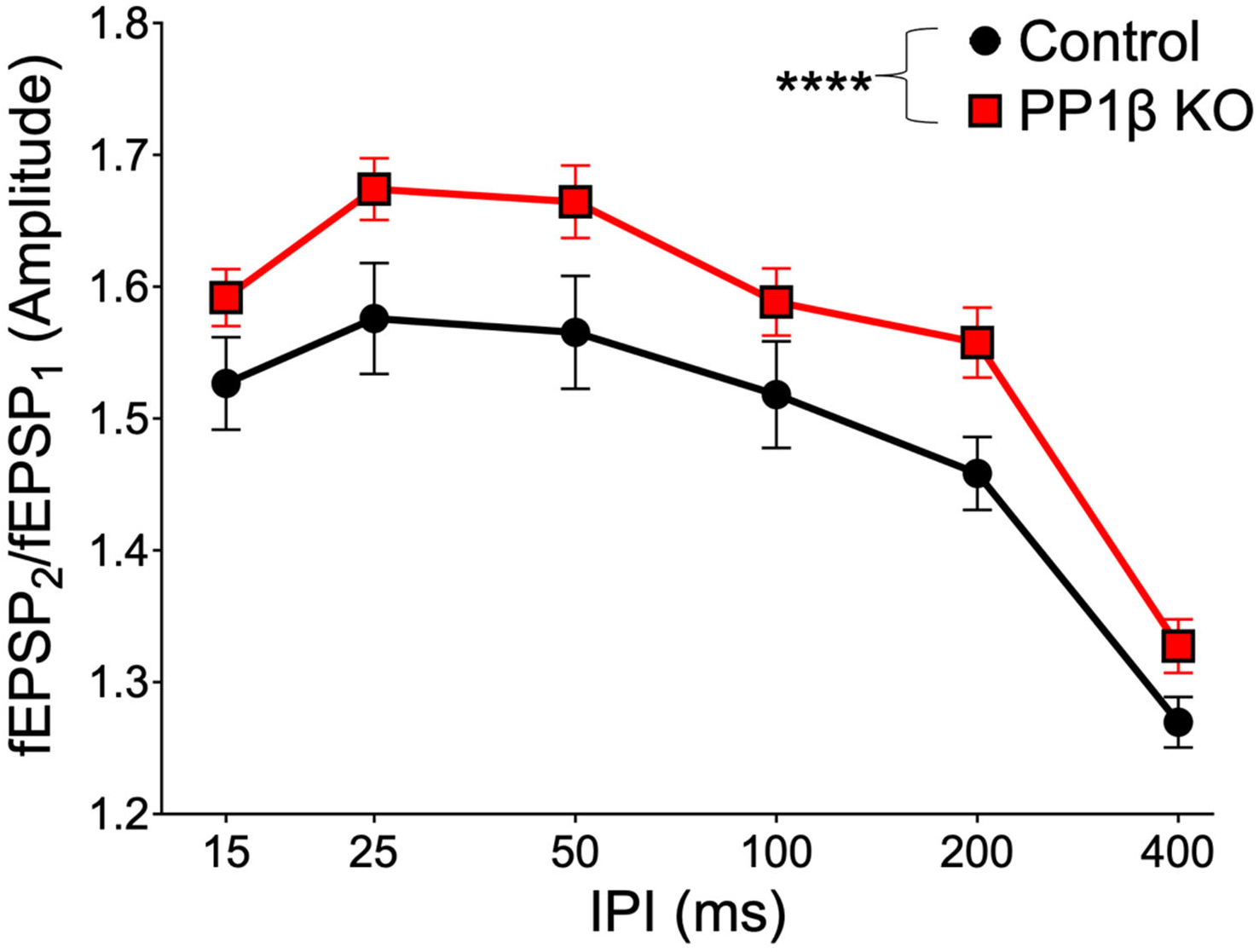
Increased paired-pulse ratio in PP1β cKO hippocampal slices. Paired-pulse ratio measured at Sch-CA1 at different interpulse-intervals (IPI) (two-way ANOVA \*\*\*\**p* < 0.0001 by genotype; n = 3 slices per animal; 3 animals per group, P10-12). Error bars denote mean +/-SEM.

Neuronal activity has been shown to positively regulate callosal myelination (Mitew et al., 2018). Decreased glutamate release in PP1β cKO mice would indicate reduced neuronal activity. We thus sought to examine the effect this would have on developmental myelination in PP1β cKO mice. We performed western blotting on cortical and hippocampal tissues and found a significant deficit in myelin basic protein (MBP) protein level in PP1β cKO cortex compared to littermate controls (Fig. 3A). There was a trending decrease in MBP that was observed in the hippocampus, but this failed to reach statistical significance (Fig. 3B).

**Figure 3.**
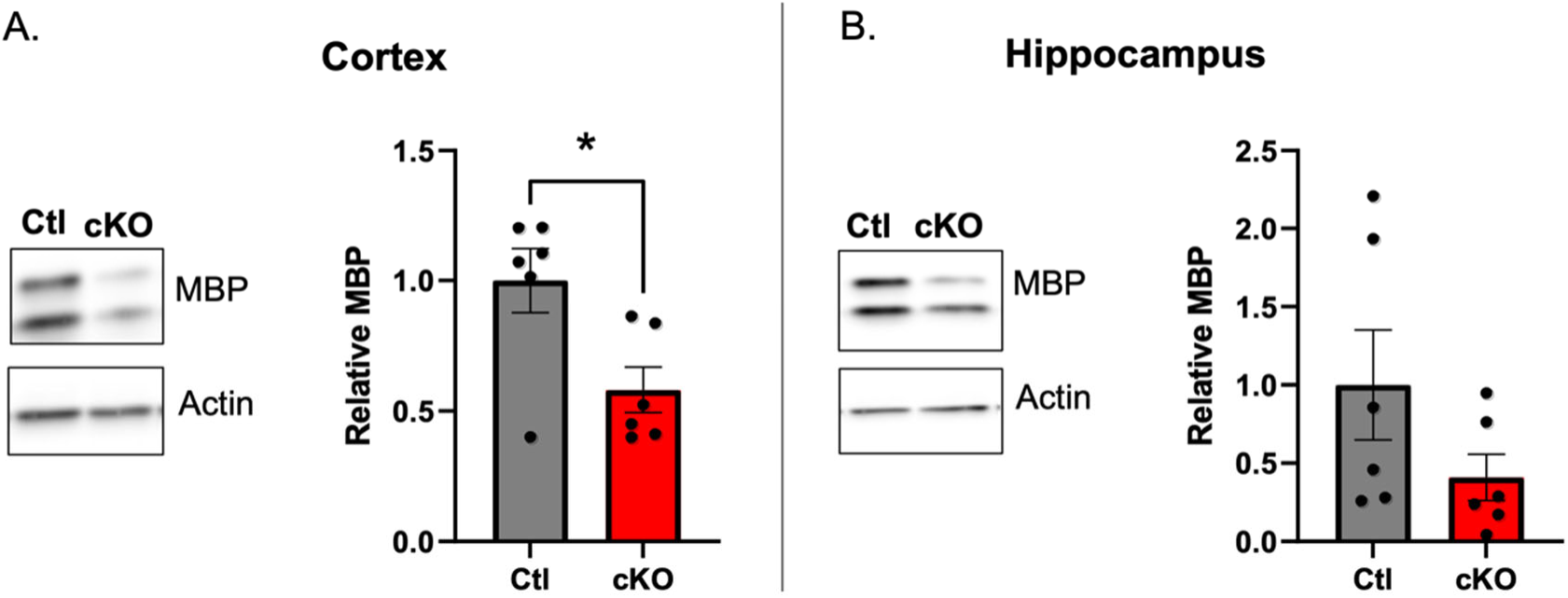
Reduced myelin basic protein expression in cortex of PP1β cKO mice. **A-B.** Western blot of cortical (**A**) and hippocampal (**B**) tissue from Ctl and PP1β cKO mice probed with antibodies against myelin basic protein (MBP) and actin (left panels). Quantification of MBP normalized to actin (right panels). **A**. two-tailed Mann-Whitney U test \**p* < 0.05. **B**. two-tailed unpaired t-test p = 0.15. n = 6 animals, P12; Error bars denote mean +/-SEM.

We next examined myelination within the optic nerve, a classic model for studying developmental myelination. In the optic nerve, myelination begins at ∼P7 and is believed to be predominantly influenced by axon diameter and not neuronal activity (Mayoral et al., 2018). We isolated these nerves from P13-14 mice and recorded compound action potentials (CAPs) to assess the electrical functionality of the axons. CAPs recorded from nerves where myelin is still developing exhibit 2 distinct peaks. The first, more rapid peak, is considered to result from action potentials propagating down myelinated axons, while the second, slower peak represents propagation down un/pre-myelinated fibers (Fig. 4A). Thus, the ratio of amplitudes of these peaks represents a measure of the relative level of functional myelin. In extracellular recordings, the amplitudes of individual peaks represent only an approximate measure of fiber density. PP1β cKO nerves had smaller peak 1 amplitudes compared to controls, with no clear difference present in peak 2 amplitudes (Fig. 4B-C). More importantly, the ratio of peak1/peak2 was significantly smaller in the PP1β cKO (Fig. 4D). No change in the velocity of either of these peaks was observed (Fig. 4E-F). Our data thus suggest a deficit in the relative proportion of functional myelinated axons in PP1β cKO nerves. Surprisingly, however, structural analysis using electron microscopy failed to detect a difference in the proportion of myelinated axons (Fig. 5A-B). Correspondingly, measurements of MBP staining density and intensity in longitudinal sections of optic nerves also failed to detect a difference between control and PP1β cKO mice (Supplemental Figure 2). Moreover, G-ratios were not significantly different between control and PP1β cKO mice, suggesting no apparent difference in the level of myelination in those axons that were ensheathed (Fig. 5C). Additionally, while the average axon diameter for each genotype did not significantly differ, the cumulative frequency distribution of axon diameters was significantly different (Fig. 5D). The cumulative distribution of axon diameters in PP1β cKO mice exhibited a shift towards smaller diameters, suggesting that a greater proportion of axons have smaller diameters in PP1β cKO optic nerves compared to controls. We did not observe a change in cell bodies in the retina. Retinal flat mounts and sections failed to detect aberrations in retinal ganglions cell (RGC) density/size, or abundance of additional support cells within the retina (Supplemental Figure 3-4). Our data thus suggest a specific role of PP1β in the nerves.

**Figure 4.**
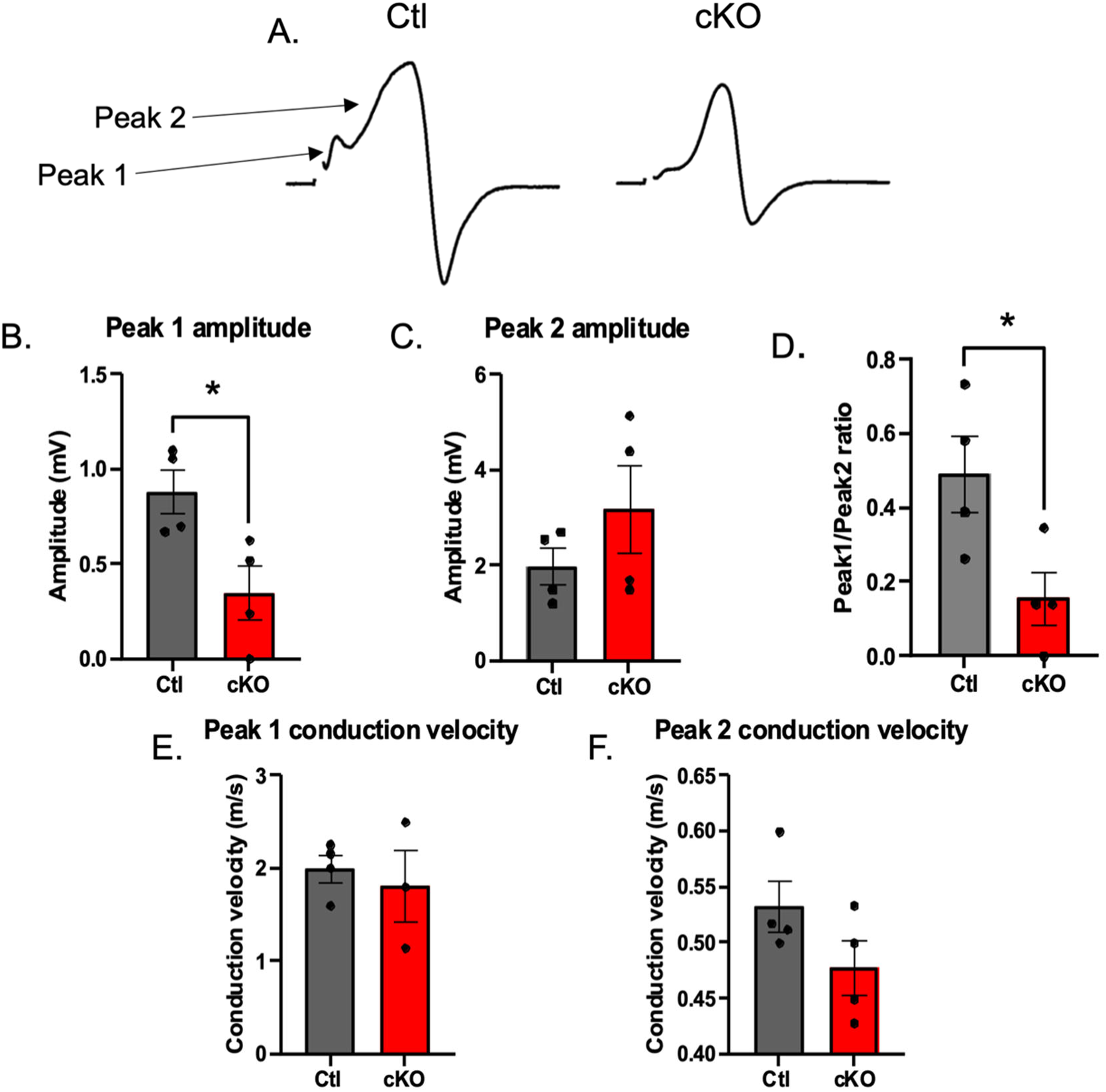
Reduced peak 1 amplitudes in optic nerves of PP1β cKO mice. **A**. Representative traces of compound action potentials (CAPs) recorded from the optic nerve (stimulation artifact removed for clarity). **B**. Measured peak 1 amplitudes (two-tailed unpaired t-test \**p* < 0.05). **C.** Measured peak 2 amplitudes (two-tailed unpaired t-test *p* = 0.28) **D**. Quantification of ratios of Peak 1 / Peak 2 CAP amplitudes (two-tailed unpaired t-test \**p* < 0.05). **E-F**. Quantification of conduction velocities from peak 1 and peak 2 respectively (two-tailed unpaired t-test *p* = 0.62, 0.15 respectively) n = 4 nerves, P13-14; Error bars denote mean +/-SEM.

**Figure 5.**
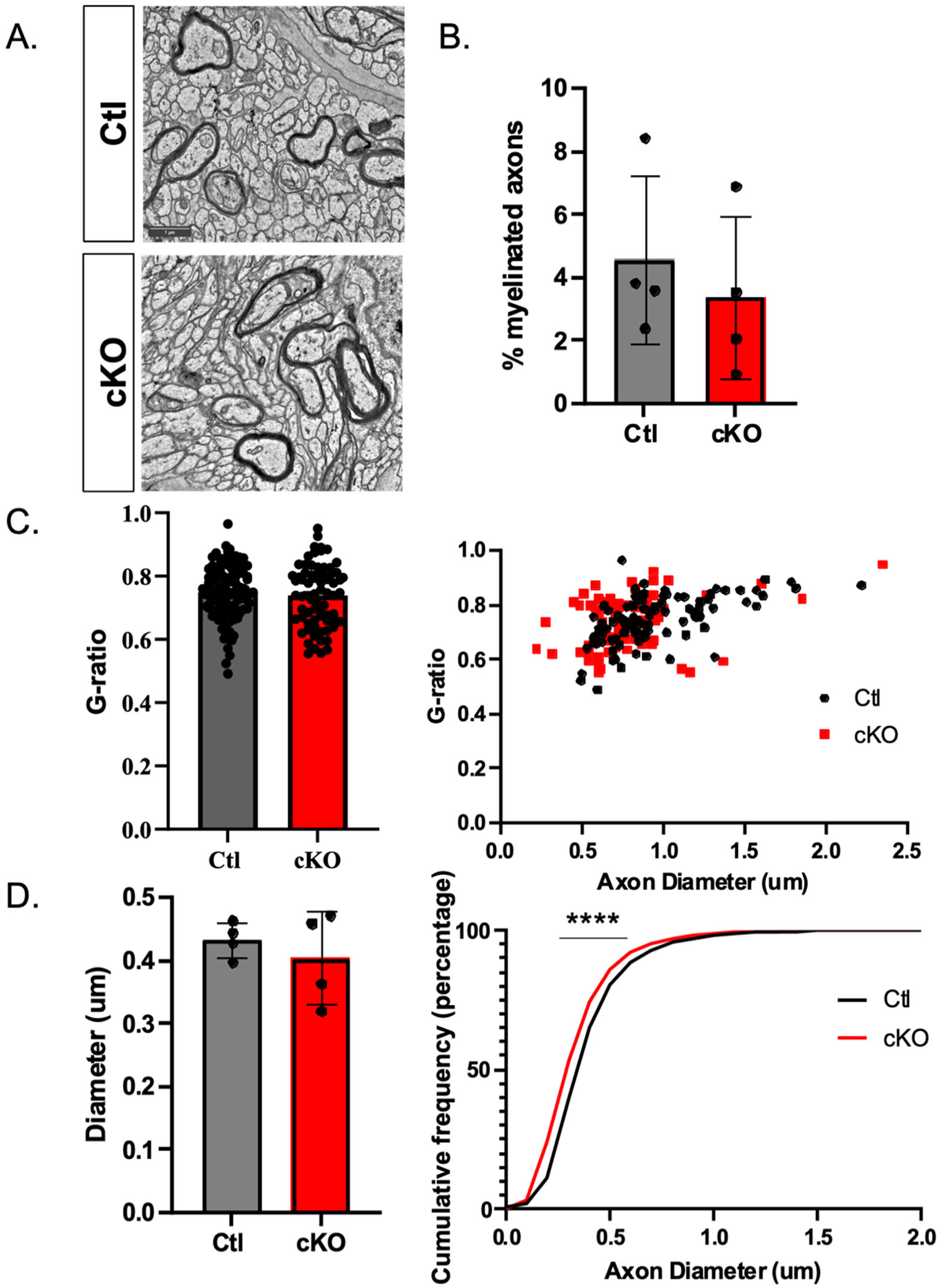
No change in myelination but a shift towards smaller axons within the optic nerve. **A.** Representative electron micrographs of optic nerve cross sections. **B**. Quantification of the percentage of myelinated axons (two-tailed unpaired t-test \**p* > 0.05 n = 4 animals); **C**. Average G-ratio of myelinated axons (left). G-ratio as a function of axon diameter (right) (two-tailed unpaired t-test *p* > 0.05; n = 4 animals; 99 ctl axons; 71 cKO axons) **D**. Average diameter of axons (left) (two-tailed unpaired t-test *p* > 0.05; n = 4 animals) with cumulative frequency distribution (right) (Kolmogorov-Smirnov test \*\*\*\**p* < 0.0001, 1300-3000 axon diameters per animal; 4 animals per group, P11-14). Error bars denote mean +/-SEM.

We next examined the nodes of Ranvier, axonal domains known to play critical roles in action potential propagation, by staining longitudinal sections of optic nerves with a pan-NaV antibody to label nodes and anti-Caspr to label paranodes. NaV clusters that aligned with paranodal Caspr were then counted to determine the density of nodes formed along the optic nerve. The density of nodes was significantly lower in PP1β cKO mice than controls (Fig. 6A-B). This was not due to deficits in NaV localization since the percentage of Caspr pairs containing positive NaV staining was similar between groups (Fig. 6C).

**Figure 6.**
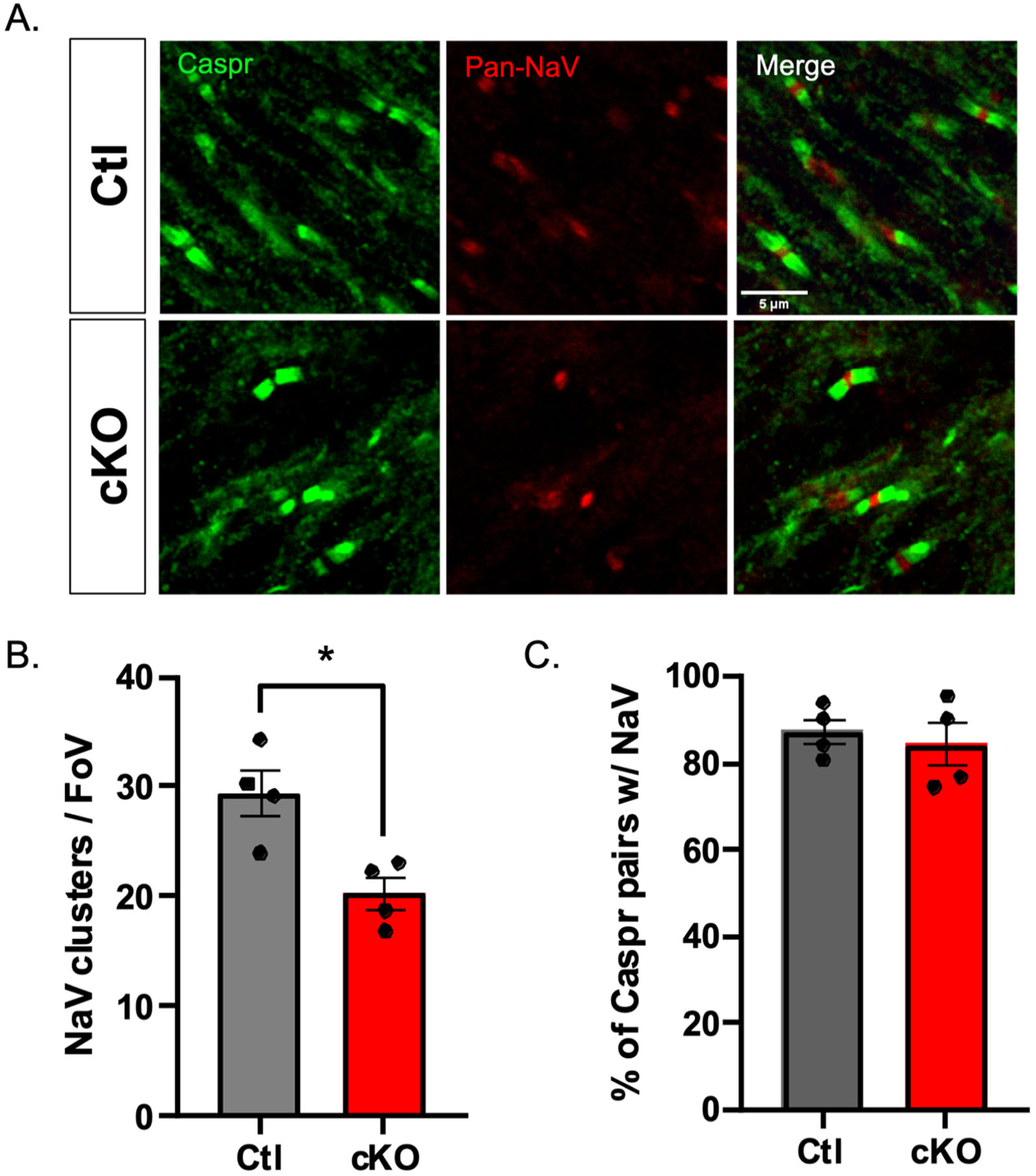
Reduced nodal NaV clusters within the optic nerves of PP1β cKO mice. **A.** Representative images of longitudinal sections of optic nerve stained with paranodal marker Caspr (green) and nodal marker pan-NaV (red). **B.** Quantification of NaV clusters associated with Caspr per field of view (FoV) **C.** Quantification of the percentage of paranodes with Caspr pairs that contain nodal NaV. (two-tailed unpaired t-test \**p* < 0.05, n = 4 animals, P14). Error bars denote mean +/-SEM.

To study the underlying mechanism of PP1β function, we probed optic nerve sections with antibodies against a known PP1β specific substrate, myosin light chain 2 (MLC2). MLC2 is a component of nodes of Ranvier and has been shown to play a role in the development of the axon initial segment, a region with structural similarity to nodes (Berger et al., 2018). Phospho-MLC2 (pMLC2) staining area normalized to that of neurofilament (NF) was found to be significantly increased in PP1β cKO optic nerves compared to control (Fig. 7A-C). Additionally, this increase in relative pMLC2 was not due to changes in axon abundance as determined by NF staining (Fig.7D).

**Figure 7.**
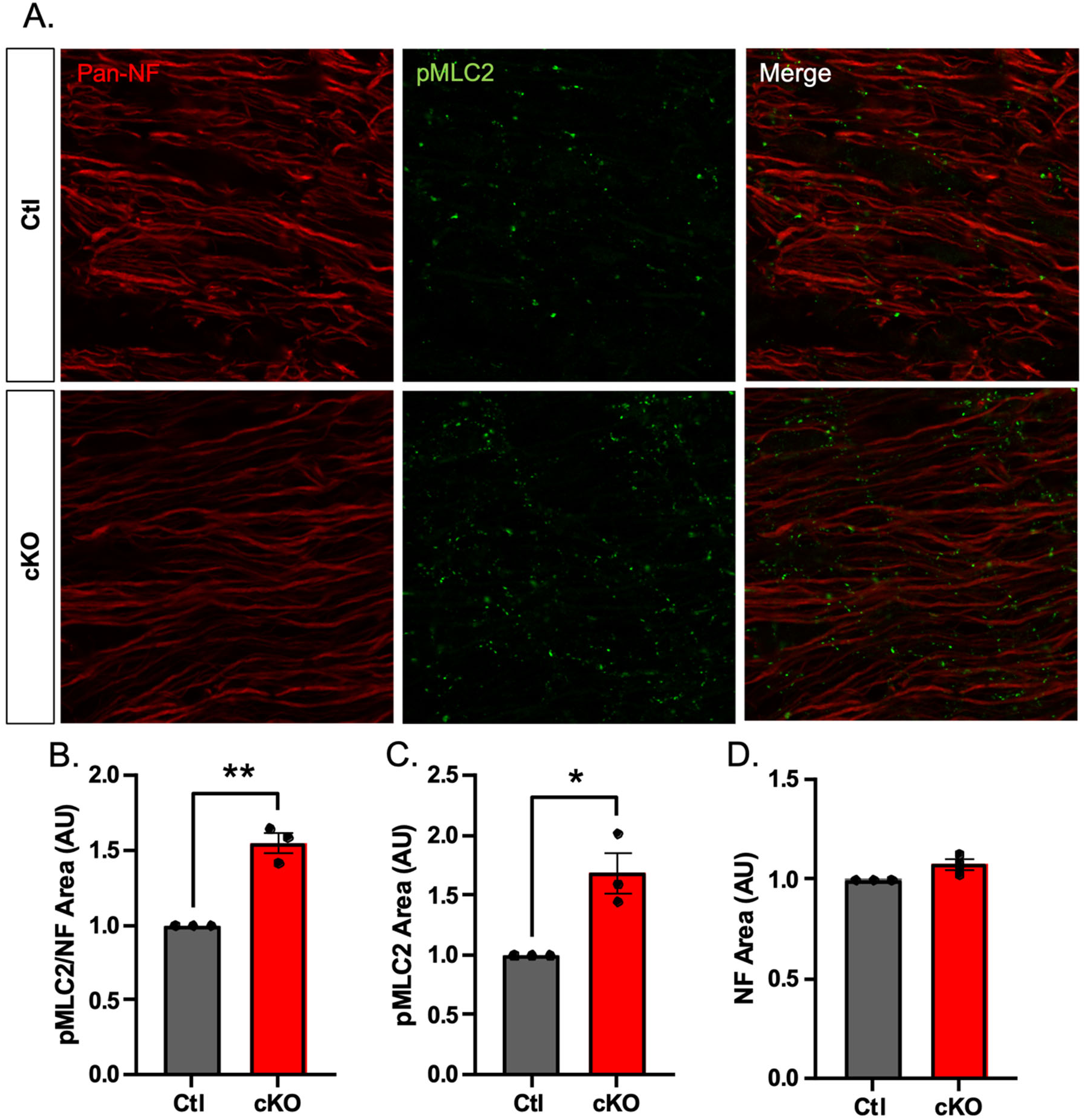
Significant increase in pMLC2 staining area in optic nerve of PP1β cKO mice. **A**. Representative images of longitudinal optic nerve sections stained with antibodies against pan-neurofilament (Pan-NF) and phospho-myosin light chain 2 (Thr18/Ser19) (pMLC2). **B**. Normalized staining area of pMLC2 relative to pan-NF (two-tailed unpaired t-test \*\**p* < 0.01). **C**. Significant increase in pMLC2 staining area (two-tailed unpaired t-test \**p* < 0.01, n = 3 animals, P13-14). **D**. No change in NF staining area. Error bars denote mean +/-SEM.

## Discussion

This work describes a number of new findings regarding the role of neuronal PP1β *in vivo*. (1) Neuronal expression of PP1β is essential for the development and survival of neonatal mice, suggesting critical roles in neuronal function. (2) There is an increase in the PPR in PP1β cKO mice at the Sch-CA1 synapses, suggesting a decrease in glutamate release. (3) We observed a decrease in MBP protein expression in cortex, suggesting a deficit in cortical myelination. (4) We observed a robust deficit in the propagation of CAP recordings in PP1β cKO optic nerve and that this is not likely due to a deficit in the frequency or thickness of optic nerve myelination. (5) We found that there is a decrease in the formation of the nodes of Ranvier, and finally (6) we showed that phosphorylated MLC2 is increased in PP1β cKO optic nerves.

To the best of our knowledge, this report is the first study to show by electrophysiological recordings a role of any PP1 isoform in presynaptic neurotransmitter release within the hippocampus. These data are consistent with previous biochemical studies using cortical synaptosomes and electrophysiological recordings of acoustic nerve synapses (Lontay et al., 2012). Our observation of reduced MBP expression in the cortex suggests a previously unknown role for PP1 in CNS myelination. A deficit in neurotransmitter release and neuronal activity in the cortex could explain the impaired MBP production that was measured by western blotting. A trending, but not statistically significant decrease in MBP in the hippocampus could suggest a different level of susceptibility to activity-dependent myelination. However, there are other possibilities that could explain this difference, including variations in recombination efficiencies or changes in neuronal activity. Furthermore, deficits in myelination were not likely a result of impaired BDNF production, a known contributor to developmental myelination (Supplemental Figure 1). However, it is possible that neuronal activity induced release of BDNF was hindered.

To determine if PP1β influenced myelination in a region that is myelinated predominately independent of neuronal activity, we examined the optic nerve. CAPs recorded from PP1β cKO optic nerves demonstrated a significant deficit in the ratio of peak 1/peak 2 amplitudes compared to littermate controls (Fig. 4). The more rapid initial peak is generally considered to be proportional to the abundance of the myelinated axons within the nerve, while the amplitude of the slower second peak is considered to reflect the relative abundance of un/pre-myelinated axons. The deficit in this ratio thus suggests that the relative functional contribution of myelinated axons is significantly reduced with neuron-specific deletion of PP1β. Interestingly, this deficit in functionally myelinated axons is also observed in nuclear inhibitor of PP1 (NIPP1) KO mice (McKee et al., 2022). However, in contrast with NIPP1 KO mice, we did not detect a significant difference in the percentage of myelinated axons in PP1β cKO optic nerves at the electron microscope level.

Additionally, while the average axon diameter did not measurably change, PP1β cKO optic nerves displayed a significant shift in the distribution of axon diameters (Fig. 5C). This increase in the relative proportion of smaller axons in PP1β cKO optic nerves coincides with previous studies investigating the role of PP1β mediated contraction of actin-cytoskeletal networks (Costa et al., 2020; Fan et al., 2017). However, it appears that this shift was insufficient to significantly impact the proportion of myelinated axons.

The decrease in peak 1/peak 2 amplitude ratios, with no accompanying change in the proportion of myelinated axons in PP1β cKO optic nerves, was unexpected, and led us to examine the possibility of impaired formation of nodes of Ranvier. The deficit in nodes that was found can potentially explain the abrogated peak 1 CAP amplitudes through a failure of action potential propagation. The mechanism linking PP1β to node formation is likely via MLC2, a PP1β specific substrate present at nodes of Ranvier (Berger et al., 2018). While the role of MLC2 in nodes of Ranvier is currently unknown, previous studies have suggested it is important in the formation of the axon initial segment (AIS) and in modulating axon diameter (Berger et al., 2018; Costa et al., 2020; Fan et al., 2017). The AIS is considered to be structurally related to nodes of Ranvier due to the abundance of shared scaffolding proteins critical for structural integrity (Rasband & Peles, 2021). We show here that PP1β deletion leads to both elevated phosphorylation of MLC2, and also impaired node formation, suggesting a possible causal link between the two. Future work will seek to delineate the specific mechanisms by which PP1β and MLC2 modulate formation of nodes of Ranvier.

It is useful to compare these results with those in NIPP1 deficient mice (McKee et al., 2022). Mice in which NIPP1 is conditionally deleted from neural precursor cells exhibit deficits in cortical MBP expression, and reduced CAP peak 1/peak 2 ratios. EM studies uncovered a significant reduction in the percentage of myelinated axons in the NIPP1 cKO optic nerve. This difference suggests a non-neuronal role of PP1 in developmental myelination. Additionally, NIPP1 binds all PP1 isoforms, and its neuronal deletion could thus affect them all. PP1β cKO mice may exhibit some level of compensation from PP1α or PP1γ. PP1α and PP1γ null mice thus need to be generated for addressing this question in the future.

This study emphasizes the critical role neuronal PP1β plays in CNS function and development. Future investigations will attempt to uncover PP1β specific interacting proteins responsible for regulating PP1β activity in these cellular mechanisms (Foley et al., 2023). Our study will also assist to determine the mechanism by which PP1β *de novo* mutations lead to intellectual developmental disorders in human patients (Gripp et al., 2016; Hamdan et al., 2014; Ma et al., 2016) .

## Acknowledgements

This work was supported by NIH R01MH128279A1 to HX, EY018606 and an unrestricted grant from the Research to Prevent Blindness to the Department of Ophthalmology at the University of Rochester to RL, and the URMC Del Monte Neuroscience Institute Schmitt Pilot Program in Integrative Neuroscience to PS and HX. The authors declare no competing financial interests.

## Supplemental Figures

**Figure S1.**
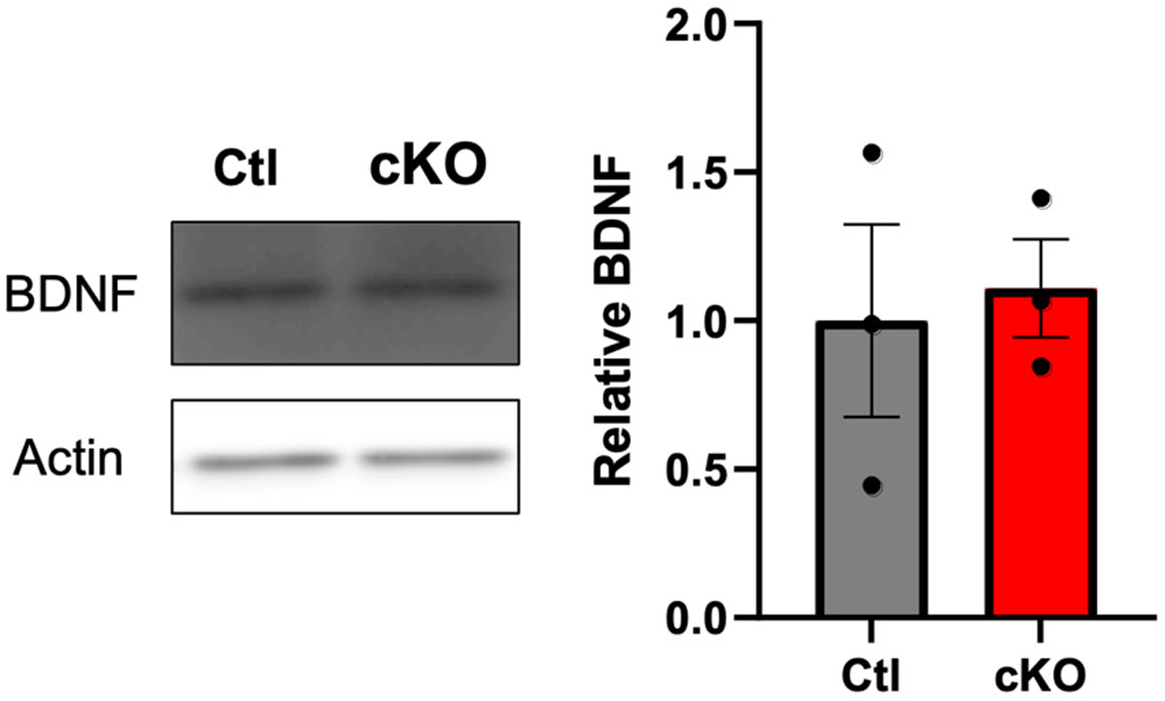
No change in BDNF in PP1β cKO cortex. Representative western blot probed with antibody against active BDNF (left) with quantification normalized to actin (right) (n = 3); Error bars denote mean +/-SEM.

**Figure S2.**
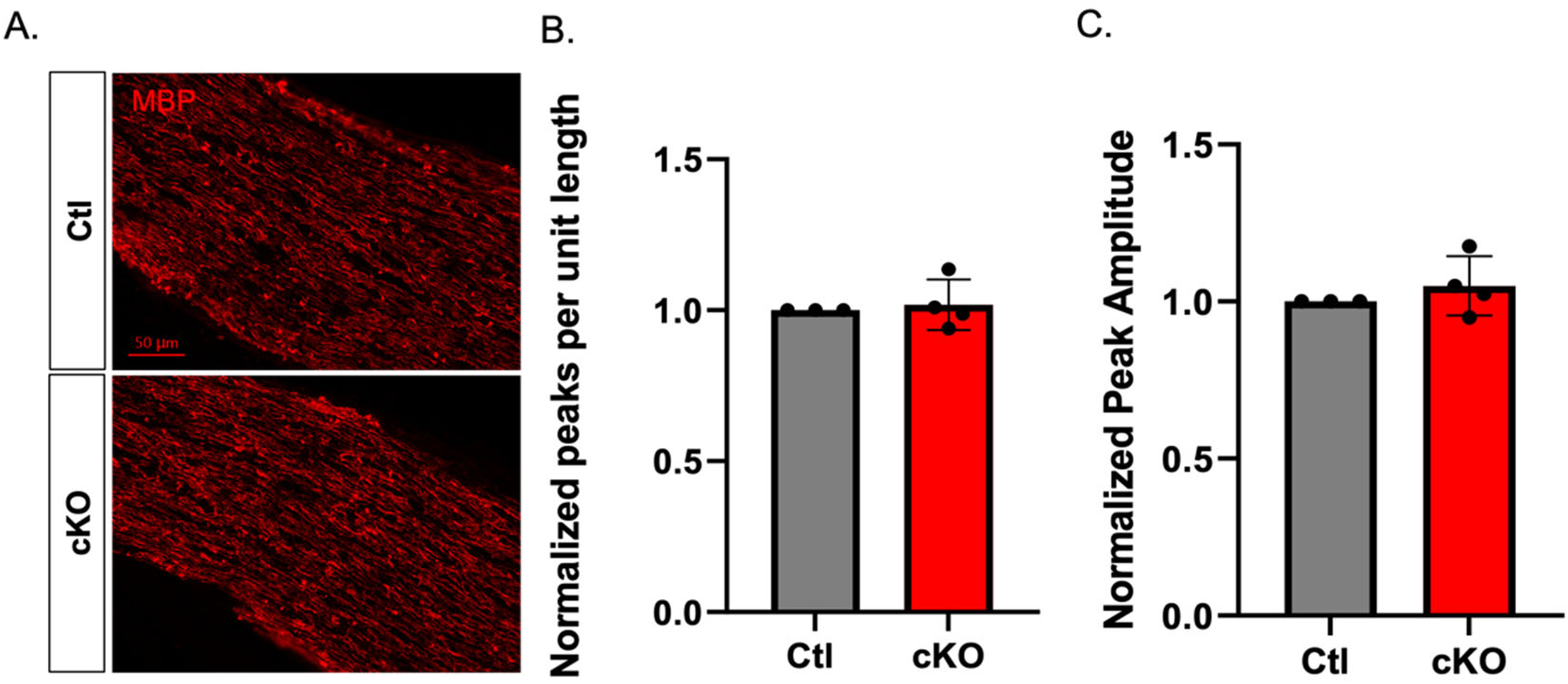
No change in MBP staining density or intensity observed in PP1β cKO optic nerves. **A**. Representative images of longitudinal optic nerve sections stained for myelin basic protein (MBP). **B-C**. Quantitation of the density (**B**) and amplitude (**C**) of pixel intensity peaks averaged across 6 linear measurements per image normalized to control (n = 3-4 animals); Error bars denote mean +/-SEM.

**Figure S3.**
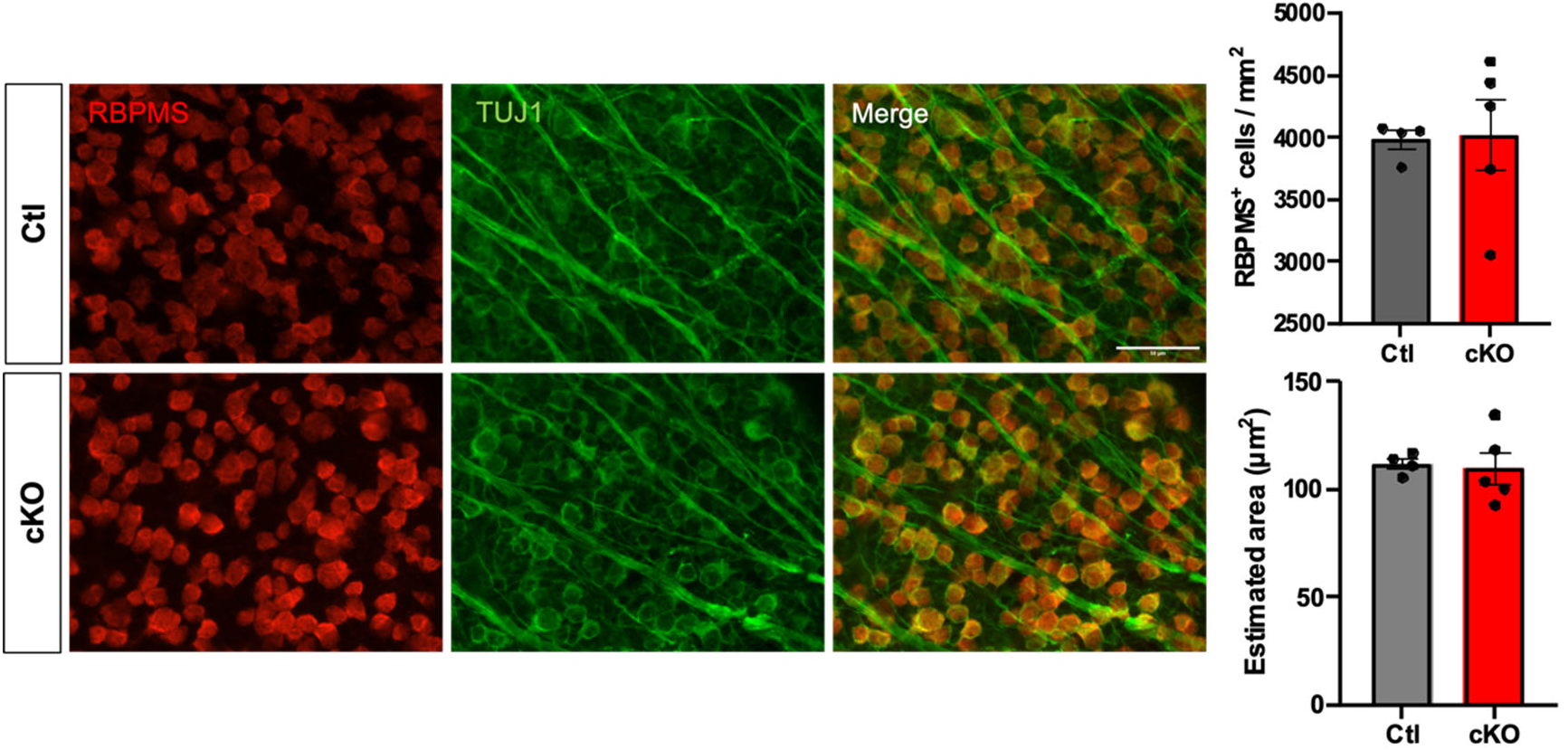
No difference in RGC density or size in PP1β retina. **A.** Representative images of retinal flat mounts stained with antibodies against RNA binding protein, mRNA processing factor (RBPMS) to label RGCs and Class II beta tubulin (TUJ1) to label axons. **B**. Quantification of RGC density. **C**. Average RGC size was estimated by dividing the area of positive RBPMS staining by the number of cells counted per image (n = 4-5 eyes per genotype). Error bars denote mean +/-SEM.

**Figure S4.**
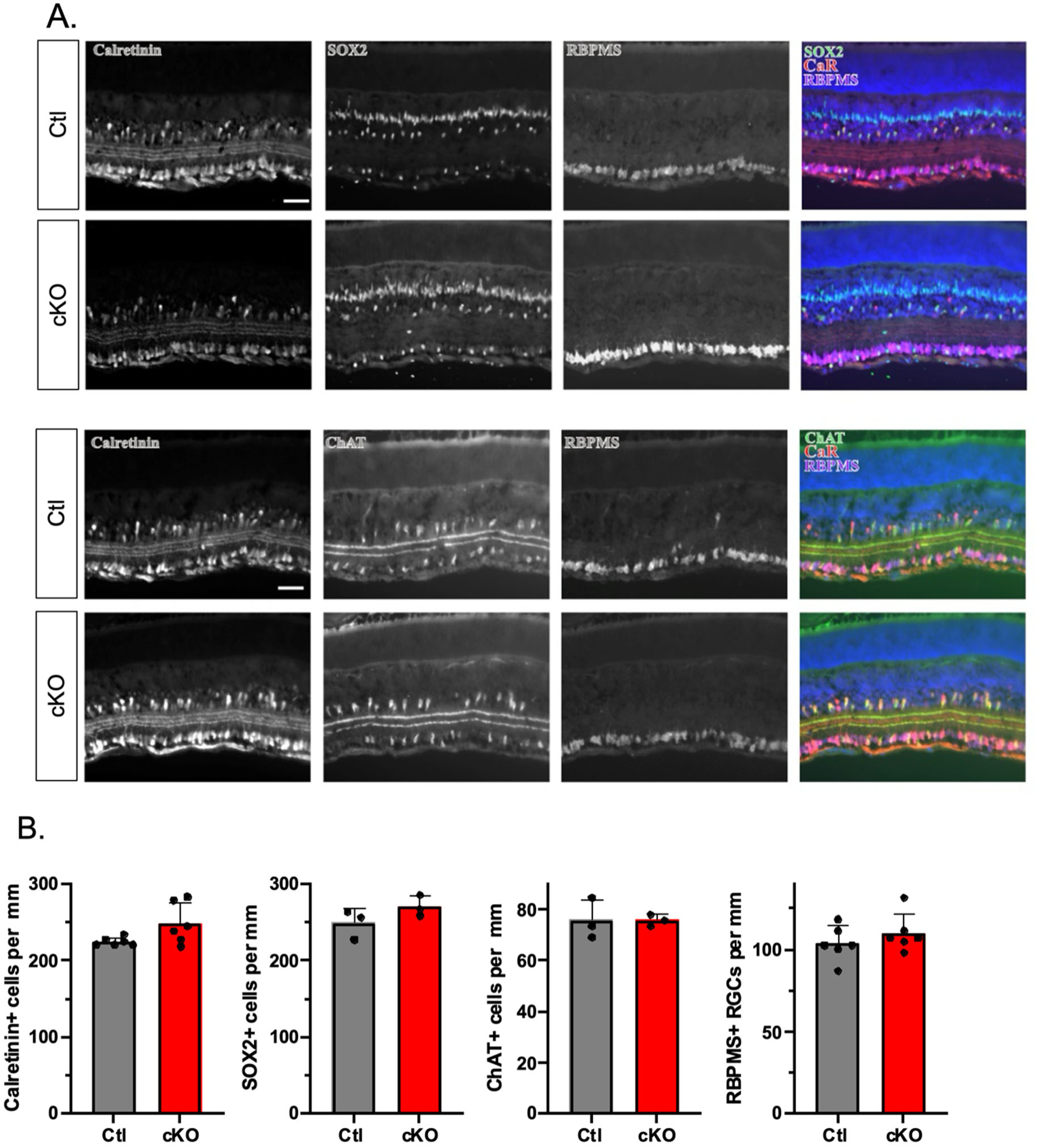
No difference in retinal neuron subtypes present in PP1β cKO mice. Representative images of frozen sections of retinas assessed for differences in subsets of amacrine cells (ChAT, calretinin, SOX2), Muller glia (SOX2), and RGCs (RBPMS, calretinin). **A.** Control and PP1β deficient animals showed similar levels of staining for calretinin, SOX2, RBPMS, and ChAT in adjacent sections. **B**. Quantification of calretinin+, SOX2+, ChAT+, and RBPMS+ cells per mm showed no significant differences between control and cKO animals (ns, p>0.05, two-tailed unpaired t-test, n≥3 sections per group). Error bars denote mean +/-SEM. Scale bar = 50µm.

